# PhageHost-GNN: novelty-aware and locally adaptable strain-level *Klebsiella*-phage prediction

**DOI:** 10.64898/2026.10.07.757501

**Authors:** Biffon Manyura Momanyi, Kunhao Qin, Kai Zhou, Lu Fan

## Abstract

Interest in bacteriophages as alternative treatments for bacterial infections has grown amid the ongoing crisis of multidrug resistance. However, their narrow host range makes it labor-intensive to identify matching phages for bacterial isolates using conventional experimental methods. Although existing computational models based on biomolecular features have shown promise for phage-host matching, the performance of alternative model settings, including data embeddings, across datasets with different degrees of novelty remains underexplored. This study introduces PhageHost-GNN, a novelty-aware framework using routed models to predict strain-level *Klebsiella*-phage interactions, evaluated under three clinical scenarios. Using multiple embedding models to encode sequence, structural, and genomic-context features, the framework achieved AUPRCs reaching 0.784, 0.731, and 0.642 in host-unseen, phage-unseen, and both-unseen *in silico* evaluations, respectively. Benchmarks on experimentally validated datasets showed the clearest advantage over fixed embeddings when both hosts and phages were new to the trained model. In addition to direct inference and *de novo* training, the framework further implements a fine-tuning pipeline to adapt existing models to local custom datasets while reducing model-training time. Overall, PhageHost-GNN provides a flexible framework for model selection, evaluation, and local adaptation in practical phage-therapy screening.

**New aspects/highlights:**

- Novelty-aware and multi-model routing for phage-host prediction.
- Implementation of three clinically practical inference settings.
- Flexible clinical adaptation via *de novo* training or fine-tuning based on the novelty of local strain datasets.

## Introduction

The growing use and misuse of antibiotics have accelerated antibiotic resistance, increasing the need for bacteriophage (phage)-based treatments to combat antibiotic-resistant bacterial pathogens in routine clinical practice. As phages often have a highly specific, strain-level host range, it is essential to identify productive interactions between candidate phages and bacterial hosts for the successful application of phage therapy [1, 2]. In routine practice, clinical laboratories often maintain a local phage library, a collection of phages isolated from environmental sources, and build an infection spectrum by conducting assays, such as spot/plaque assays and microfluidic polymerase chain reaction, to test interactions between stock phages and bacterial strains [3, 4]. However, these experiments are expensive, labor-intensive, and time-consuming, especially for large-scale applications [3, 5, 6]. This can delay the identification of sufficiently active phages for inclusion in phage cocktails against newly isolated pathogenic strains.

In recent years, *in silico* prescreening approaches have been developed to accelerate the identification of active phages. Bioinformatics- and machine-learning-driven methods have been applied to predict strain-level matches between phages and bacteria [7]. These studies often utilize features of phage receptor-binding proteins (RBPs, e.g., tail fiber proteins), host receptor proteins, or proteins that synthesize host receptor molecules (e.g., lipopolysaccharide and capsular polysaccharide), and train models on experimentally verified phage-host pairs to predict potential phages for new hosts or hosts for new phages. In recent studies, language-model-based and three-dimensional structure-based embeddings have been applied to capture features of phage-host interaction proteins [8–10]. While these models have achieved promising predictive performance for clinical applications, they have important limitations. First, previous studies either utilized one specific algorithm to embed features of phage or host proteins [2, 8, 11] or concatenated/fused multiple embeddings [12]. Few have tested model performance and generalizability for phage or host entities with different degrees of novelty and subsequently developed processes to optimize model selection based on the novelty of query entities. Second, published models were often trained on specific datasets generated in a single laboratory or hospital, which were often limited and geographically biased, considering the vast global diversity of bacterial and phage strains [13]. Accordingly, performance may decline when such models are applied to strains isolated in other laboratories or hospitals, especially in geographically distant regions [14]. Therefore, there is a need for a scalable tool that can incorporate newly published or clinic-specific local datasets (i.e., genomes or key proteins of newly isolated phage or host strains, and new experimentally verified interactions) to continually improve its prediction accuracy. Users may also need to adapt the published models to their local phage/host data for clinic-specific phage-therapy workflows.

In this study, we develop a novelty-aware framework named PhageHost-GNN, which supports local adaptation to custom datasets to predict strain-level phage-host interactions. The framework uses a GNN-based deep learning approach; the graph represents the phage-host infection spectrum and is designed to incorporate new phage and host strains as additional datasets become available. It allows users to test the performance of different embedding methods for protein sequences and select the ones best suited to phage or host entities at different levels of novelty relative to the training dataset to improve the model’s predictive performance. A fine-tuning pathway enables efficient addition of new datasets to adapt the model to local prediction needs. We used recently published interaction datasets of *K. pneumoniae* and phage strains to demonstrate the framework’s implementation. Benchmarking of our model-routing setting with fixed embedding strategies shows competitive host-unseen performance, with larger gains under the both-unseen setting, where both phages and hosts are novel to the trained model. The implementation of this framework is publicly available on GitHub, supporting customization for practical phage-therapy screening.

## Results

### Overall design and implementation of PhageHost-GNN

Feature extraction and embedding. During adsorption, many *Klebsiella* phage RBPs recognize capsule-associated structures, making capsular polysaccharide an important host-range determinant [15, 16]. Many characterized *Klebsiella* phages target a single capsule type, and alterations in capsule production or capsule-associated structures can confer resistance [17, 18]. In the context of protein language representations and structural conservation [9, 13], phage RBPs and bacterial K-locus proteins are extracted from the respective genomes and three feature types—protein sequence, protein structure, and genomic context—are represented using five embedding strategies, including ESMC (an ESM-family pretrained protein language model whose representations can encode contact and structural information without direct structural input) [19, 20], ESM2 (a pretrained protein language model) [21], LucaOne (multi-modal model pretrained on gene sequences and annotations, and protein sequences, structures and annotations) [22], AlphaFold 3 (AF3, used to predict protein three-dimensional structures) [23, 24], and BacFormer (pretrained on genomic context) [25] (Fig. 1A). For BacFormer, ESM2 protein embeddings were contextualized using gene order determined through Prodigal-predicted gene coordinates, thereby capturing genomic-context information from surrounding proteins for final protein embeddings. Numerical vectors are generated for the four language model embeddings, while AF3-based structural embeddings are extracted by the ESM-IF1 encoder [26]. Because each phage contains one or more RBPs and each bacterium contains multiple K-locus proteins, the corresponding vectors are aggregated by a column-wise mean into one vector per phage or bacterium.

**Fig. 1:**
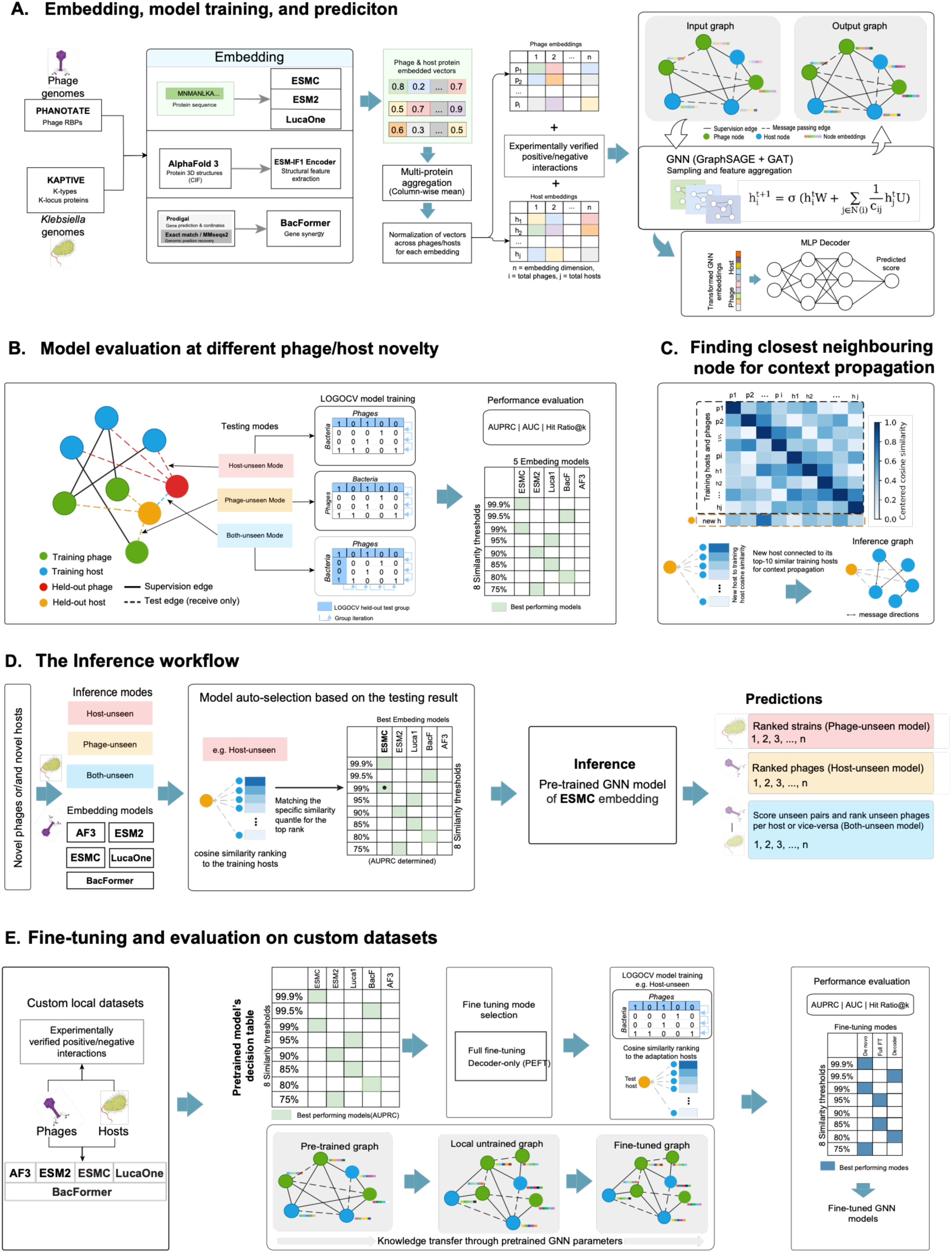
An overview of the proposed PhageHost-GNN framework. **(A)** Embedding, model training, and prediction. Feature extraction and embedding of phage RBPs and bacterial K-locus proteins are performed using pretrained language models and AF3. Graph inductive representation learning is followed by an MLP decoder that produces ranked predictions. **(B)** Model LOGOCV evaluation at different phage/host novelties. Three inference modes are evaluated. **(C)** Nearest-neighbour context propagation. **(D)** The inference pipeline, which first determines the similarity between query hosts or phages to the training dataset, and automatically assigns them to the corresponding model quantiles for inference. **(E)** Fine-tuning pipeline for local adaptation on custom datasets, allowing users to produce prioritized phage-host interactions for subsequent experimental validation.

### Model training

The proposed GNN framework employs GraphSAGE for neighborhood sampling and aggregation and GATv2 for dynamic attention, using multi-instance language-model and structure-derived embeddings as node features (Fig. 1A). Phages and hosts are represented as distinct nodes, with aggregated RBP and K-locus vectors as their initial features and experimentally verified interactions as supervision edges. Top-10 nearest-neighbor phage-phage and host-host cosine-similarity edges provide graph context through message passing. This design supports local adaptation by allowing user-specific interaction data to update phage-host representations while retaining context learned from the pretraining dataset.

### Inference modes

Model prediction is configured in three modes representing three practical settings. Specifically, the ‘host-unseen’ mode mimics a scenario where new *Klebsiella* strains are isolated in clinical practice, and *in silico* prescreening of a local phage library for active phages is required (Fig. 1B). The ‘phage-unseen’ mode mimics a scenario in which new phages are isolated from the environment, and *in silico* prescreening for active interactions with known *Klebsiella* strains is required. Finally, the ‘both-unseen’ mode represents a clinical scenario in which both query phages and bacterial hosts are new to the training dataset. During inference, new entities are connected to their 10 most similar entities in the training graph based on cosine similarity between standardized embeddings (kNN auxiliary edges) through receive-only connections, allowing them to receive additional graph context without contributing to the training graph (Fig. 1C). Specifically, in the host-unseen mode, new hosts are connected to their top-10 training hosts, in the phage-unseen mode, new phages are connected to the top-10 training phages, while in the both-unseen mode, new phages and new hosts are both connected to their respective top-10 training phages and hosts to facilitate graph context propagation before final prediction by the multilayer perceptron (MLP) decoder.

### Model evaluation

Predictive performance and generalizability of models for phage-host matching are evaluated under the above-mentioned three modes (Fig. 1B). To minimize data leakage in model evaluation, bacterial and phage strains are clustered, using thresholds defined by a series of quantiles (from 99.9% to 75%) based on protein embedding cosine similarities under the leave-one-group-out cross-validation (LOGOCV) approach, in which one group is iteratively held out from the training graph and used to evaluate generalization on previously unseen entities. Entity grouping was performed before LOGOCV to determine held-out groups, after which train-independent feature processing (including feature normalization), kNN graph construction, and model fitting were based on training fold entities, while the test fold entities were used for model evaluation.

Phage or host entities of the held-out cluster are projected to the respective model embedding spaces and compared to training entities using centered cosine similarity (Fig. 1C). For evaluation, the AUPRC, AUROC, and hit ratio at K (HR@K) are calculated. While the commonly used AUROC measures the ability of a model to rank positive interactions above negatives, it may provide an overly optimistic assessment of practical retrieval performance in highly imbalanced datasets [27]. Because the false-positive rate is normalized by the total negative instances, AUROC can remain high even when the absolute number of false positives is substantial. Instead, AUPRC summarizes the precision-recall trade-off across prediction thresholds and is particularly informative when positive interactions are rare [47, 48], whereas the HR@K measures the proportion of eligible queries (those with at least one experimentally verified interaction) for which at least one true match is retrieved among the top-K predictions, thereby reflecting the model’s practical top-ranking performance. Therefore, AUPRC is used as the main performance evaluation metric, followed by AUROC and HR@K, while sensitivity and specificity are also used at inference to evaluate binary classification from complementary perspectives under positive and negative class predictions.

### The inference workflow for new entities by model selection

New entities are embedded using the five methods, and the resulting vectors are projected to the respective model embedding spaces and compared to training entities to identify the corresponding similarity-quantile assignments for the new entities (Fig. 1D). The same approach is independently employed for phages and hosts in both-unseen inference, with new phages and new hosts routed to different similarity quantiles and then assigned to the lower of the two quantiles, defined as pair_q = min(host_q, phage_q), ensuring support for both entities. For every novel pair, the best trained model per assigned quantile is selected using the highest pooled LOGOCV AUPRC, with AUROC employed as a tie-breaker to ensure novel entities are evaluated using the most predictive quantile model.

### Fine-tuning function

This function allows users to adapt pretrained models to small local datasets without retraining from random initialization. In this study, local adaptation is evaluated using full fine-tuning and decoder-only fine-tuning modes (Fig. 1E). Full fine-tuning updates the graph encoder and MLP prediction decoder, whereas decoder-only fine-tuning freezes the graph encoder and updates only the MLP decoder, significantly reducing the number of trainable parameters.

### Dataset features

The Boeckaerts et al. [11] (Boeckaerts2024) dataset, which combines the Beamud et al. [1] (Beamud2023) and Ferriol-González et al. [28] (Ferriol-Gonzalez2024) datasets, and the Wu et al. [8] (Wu2026) dataset were used for model training and evaluation (Table 1). The former contains 333 positive and 9,673 negative experimentally verified interactions, whereas the latter contains 1,800 positive and 7,154 negative interactions, corresponding to positive-to-negative ratios of approximately 1:29 and 1:4, respectively. Additionally, 28 *K. pneumoniae* bacterial strains and 17 phages were obtained from the Boeckaerts et al. [11] study (Boeckaerts2024-in-vitro) for host-unseen inference and retrospective benchmarking, providing an independent evaluation of model performance on novel entities using a model-selection implementation. The Beamud2023 dataset was also used for retrospective benchmarking on models trained on the Wu2026 dataset under the both-unseen setting, and to test the fine-tuning approach on those models. The genome lists and the number of RBPs per phage and K-locus proteins for hosts are summarized in Supplementary Tables S1–S4.

**Table 1:** The datasets used in this study.

| Dataset | Utilization | Hosts | Phages | Interactions (pos, neg) | Reference |
| --- | --- | --- | --- | --- | --- |
| Wu2026 | Training and testing | 121 | 74 | 8,954 (1,800, 7,154) | [8] |
| Boeckaerts2024 (a combination of Beamud2023 and Ferriol-Gonzalez2024) | Training and testing | 200 | 105 | 10,006 (333, 9,673) | [11] |
| Boeckaerts2024-in-vitro | Host-unseen retrospective benchmarking for the models trained on Boeckaerts2024 | 28 | 17 (overlapping with Ferriol-Gonzalez2024) | 476 (33, 443) | [11] |
| Beamud2023 | Both-unseen retrospective benchmarking for the models trained on Wu2026 dataset.<br>Comparison of direct inference, fine-tuning of Wu2026-pretrained models, and *de novo* training | 138 | 46 | 6,348 (153, 6,195) | [1] |
| Ferriol-Gonzalez2024 | As part of Boeckaerts2024 | 62 | 59 | 3,658 (180, 3,478) | [28] |

### *In silico* performance evaluation on different embedding models

Across the three inference modes, performance of the Wu2026-trained models declined as held-out phages or hosts became less similar to the training entities, although fold composition also varied among embedding models and quantiles. Specifically, models performed best for held-out strains closely related to those in the training sets, i.e., from 99.9% to 99% similarity quantiles (Fig. 2 and 3). Below the 99% similarity quantile, model performance declined with decreasing similarity of the held-out strains, but the declining curves varied between embeddings. For example, in the host-unseen mode, while the AUPRC scores of the other three models declined from above 0.7 to around 0.5 or even 0.376 at 75% similarity, the AF3- and ESMC-embedding models maintained AUPRC values around 0.7 for more novel hosts (Fig. 2A).

**Fig. 2.**
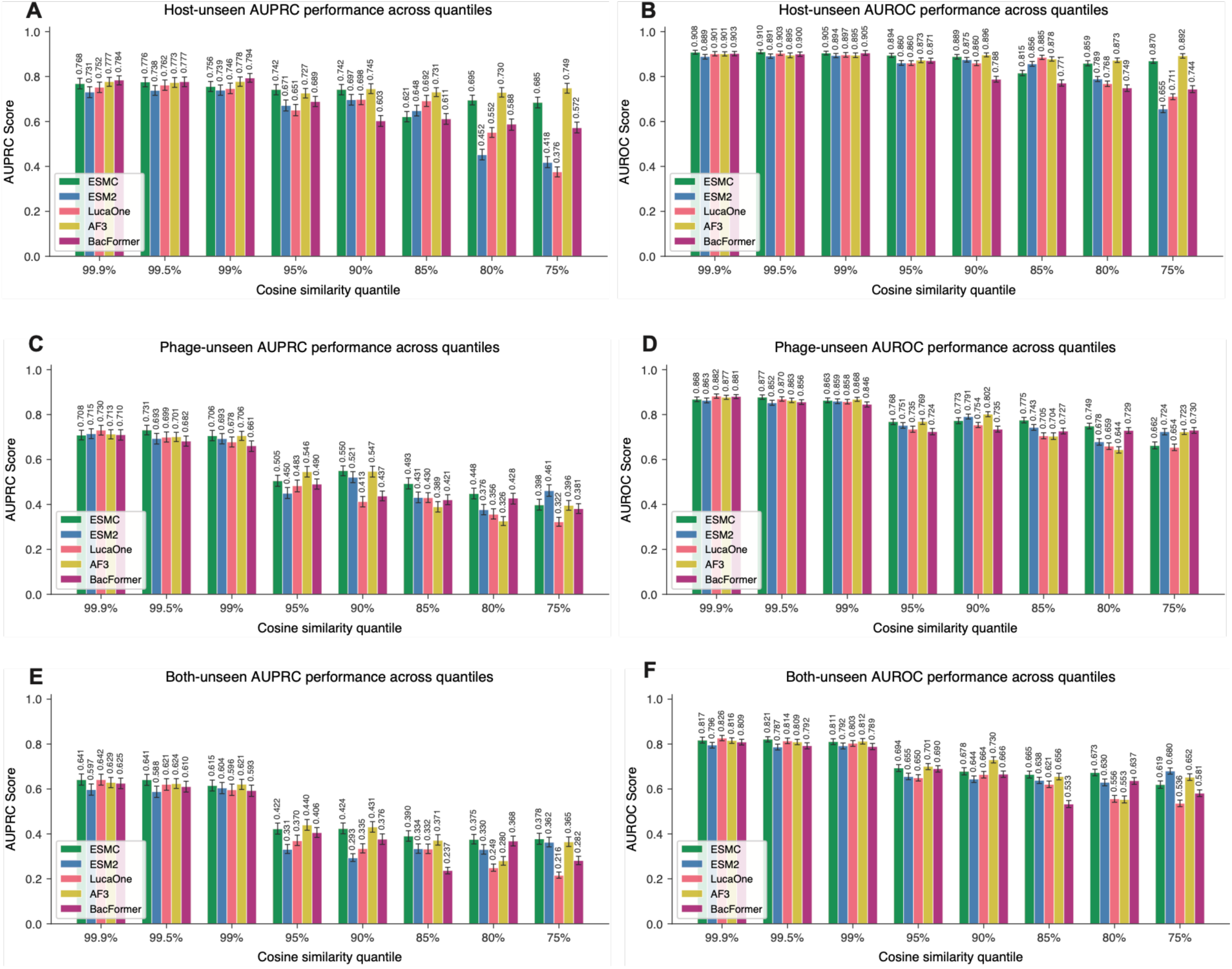
AUPRC and AUROC performance of the PhageHost-GNN models of different embeddings trained on the Wu2026 dataset on hosts/phages of different novelties. LOGOCV was conducted on the Wu2026 dataset at each cosine-similarity threshold. The held-out cluster was used for inference, while the remaining clusters were used for model training. (A, B), (C, D), (E, F) show the AUPRC and AUROC scores obtained under the host-unseen, phage-unseen, and both-unseen inference modes, respectively.

**Fig. 3.**
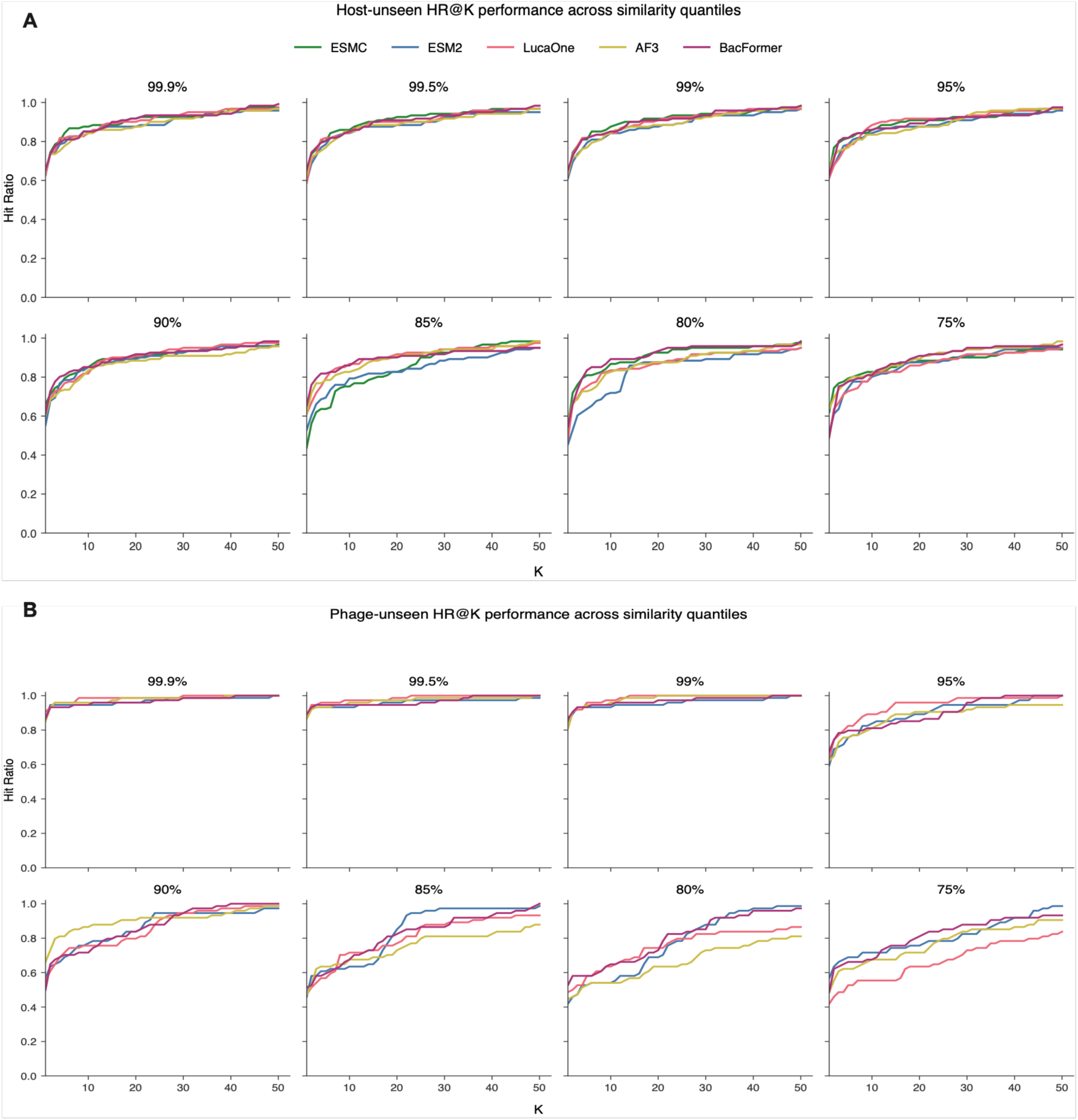
HR@1-50 curves for the PhageHost-GNN models of different embeddings trained on the Wu2026 dataset on hosts/phages of different novelties under host-unseen (A) and phage-unseen (B) inference modes. LOGOCV was conducted on the Wu2026 dataset at each cosine-similarity threshold. The held-out cluster was used for inference, while the remaining clusters were used for model training.

The best-performing embedding varied across inference modes and novelty levels (Fig. 2 and 3). For highly similar queries, differences among embeddings were generally smaller; larger differences were observed in queries of lower similarity, with no single embedding consistently outperforming the others across all phage or host similarity levels.

AUPRC values in the phage-unseen and both-unseen modes were generally lower than those in the host-unseen mode, and the performance declined more drastically after the 99% similarity quantile, possibly reflecting the challenges for models to capture highly divergent phage protein sequences and highly mosaic phage genomes (Fig. 2 and Fig. 3) [13]. Moreover, the both-unseen mode is the most challenging setting for the GNN model because of structural isolation and the two proxy paths (phage-phage and host-host), which could increase edge noise during context propagation (Fig. 1B). As expected, this mode yielded even lower AUPRC values than the phage-unseen mode (Fig. 2C, E).

Similar to AUPRC, the AUROC and HR@K metrics showed similar performance trends across decreasing phage and host similarities, with different leading models across query-entity similarity quantiles (Fig. 2B, D, F and Fig. 3). HR@K evaluation was omitted for the both-unseen setting due to sparsity of observed interactions in the held-out clusters, which increases the risk of an inflated hit ratio (mostly 100%), especially with the presence of singleton positives within the top predictions.

A supplementary analysis was conducted on the Boeckaerts2024 dataset with a similar size but a lower positive-class prevalence. While the overall model performance of all embeddings was lower than those in the Wu2026 datasets, performance generally declined for more novel entities, and the leading model varied under different similarity quantiles (Fig. S1 and Fig. S2). The AF3- and BacFormer-embedding models achieved the highest AUPRC scores for less-similar hosts from 95% to 80% and at the 75% quantile, respectively.

### External retrospective validation using novelty-dependent model selection

Because the best-performing models varied at each similarity threshold (Fig. 2 and 3), we designed and implemented a model routing strategy for inference (Fig. 1). We used the embedding model with the highest internal AUPRC at the assigned novelty quantile because it was expected to be most informative for assessing query entities with certain novelty. To quantify the benefit of this model-selection strategy over prespecified fixed-embedding models, we conducted retrospective benchmarks against previously reported experimentally measured interactions (Table 1).

A host-unseen test was conducted on the Boeckaerts2024-in-vitro dataset using Boeckaerts2024-trained models (Table 1). Out of 28 novel host strains, 20 strains were assigned to the ESMC model at 99% and 99.5% similarity, while the remaining eight were assigned to the BacFormer at 95% and 99.5%, suggesting the query host set was highly similar to the training set (Fig. 4A, Supplementary Table S5). The distribution of inference scores was strongly bimodal, with most pairs assigned scores near zero (360 with scores below 0.2), while a smaller, high-confidence subset (93 with scores above 0.8) had scores near 1.0 (Fig. S3A, Supplementary Dataset 1). In this negative-dominated test dataset, the routed model demonstrated competitive performance relative to fixed-embedding models. However, the overall differences between the routed models and the competing models were small, reflecting the high similarity between the testing and the training sets, which provides only a limited test of model generalization. Specifically, the novelty-guided routed model correctly predicted 17 out of 33 positive interactions (51.5% sensitivity), and 395 out of 443 negative interactions (89.2% specificity). In comparison, the previously published PhageHostLearn model (XGBoost) [11], which used fixed ESM2 embeddings, correctly retrieved 15 positives (45.5% sensitivity) and 418 negatives (94.4% specificity), while fixed-embedding GNN models in this study showed sensitivity/specificity pairs of (60.6% and 88.0%), (45.5% and 88.7%), (6.1% and 98.9%), (48.5% and 89.6%), and (51.5% and 88.5%), for the AF3, BacFormer, ESM2, ESMC, and LucaOne fixed embedding models, respectively. Moreover, the routed model achieved a competitive AUPRC of 0.312 (approximately 0.069 for a random model) and AUROC of 0.815, comparable to the PhageHostLearn model (0.354 and 0.782, respectively) (Fig. 4B, C). Furthermore, the routed model was among the best performers from HR@2 to HR@7, mostly tied with the PhageHostLearn model, while showing higher top-*k* retrieval performance at larger values of *k*, including HR@8, HR@9, and HR@10, where it tied with the BacFormer (Fig. 4D), indicating competitive positive retrieval, especially at larger *k* values.

**Fig. 4:**
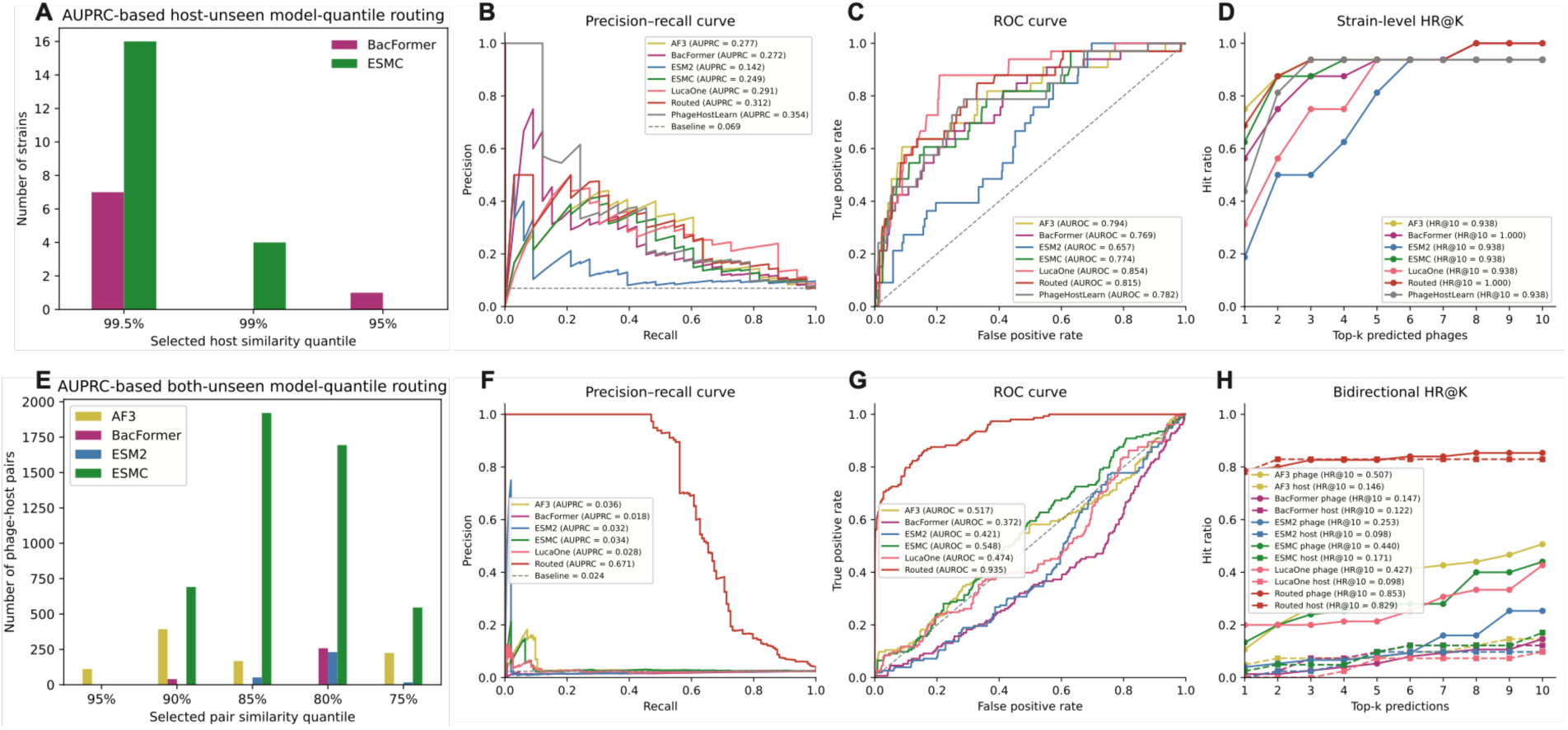
Benchmarking against experimentally measured datasets comparing similarity-guided routed models with fixed-embedding models. Inference AUPRC-determined model-quantile assignments (**A** and **E**), precision-recall curves with AUPRC scores (**B** and **F**), ROC curves with AUROC scores (**C** and **G**), and HR@K (**D** and **H**) for the Boeckaerts2024 host-unseen inference (**A-D**), and Beamud2023 both-unseen benchmarking using Wu2026-trained models (**E-H**), respectively.

The retrospective benchmark of the both-unseen setting was conducted using the China-derived Wu2026-trained models on the Spain-derived Beamud2023 dataset, which also had a low positive prevalence (2.4%). Of 6,348 novel phage-host pairs, 897, 298, 301, and 4,852 were assigned to AF3, BacFormer, ESM2, and ESMC models, respectively, across similarity quantiles (from 95% to 75%), reflecting a query dataset with low similarity to the training entities (Fig. 4E; Supplementary Table S6). Most (5,951 pairs) probability values were below 0.2, while 237 exceeded 0.8 (Fig. S3B; Supplementary Dataset 2). The routed models substantially outperformed the fixed-embedding models on this dataset in discrimination and candidate prioritization, as indicated by the large differences across the three evaluation metrics (Fig. 4F-H). At the F1-maximizing thresholds, the routed model identified 105 of 153 positives (68.6% sensitivity) and 5,854 of 6,195 negatives (94.5% specificity). The routed both-unseen model obtained an AUROC of 0.935 and AUPRC of 0.671, compared with a random AUPRC baseline of approximately 0.024. In comparison, all fixed models showed AUPRC at or below 0.036 and AUROC at or below 0.548. These models were further evaluated as a phage or a host ranking system for HR@K. As a phage-ranking system, the model correctly predicted hits for 62 out of 75 eligible hosts within the top-five predictions, corresponding to an HR@5 of 82.7%. As a host-ranking system, the model correctly predicted hits for 34 out of 41 eligible phages within the top five predictions, corresponding to an HR@5 of 82.9% (Fig. 4H).

### Fine-tuning supports transfer to local datasets

When a clinic maintains a local library of phages and host strains and seeks to build a custom model for newly isolated drug-resistant strains, it can either train a model *de novo* or fine-tune an existing model on local data. Fine-tuning may be preferable when the local dataset is small and highly imbalanced, or when computational resources are limited. To demonstrate this approach, models pretrained on the more balanced China-derived Wu2026 dataset (1,800 positives, 7,154 negatives, 1:4.0) were fine-tuned using both full fine-tuning and the decoder-only parameter-efficient fine-tuning (PEFT) on the highly imbalanced Spain-derived Beamud2023 dataset (153 positives, 6,195 negatives, 1:40.5) (Table 1). The LOGOCV approach was employed for testing on the held-out cluster under the host-unseen mode. For *de novo* training and inference, the best-performing model per quantile was retrospectively selected using the highest mean AUPRC across 10 runs, representing an oracle upper bound comparison (Fig. 5A-C, Fig. S4). For fine-tuning and inference, we used the embedding method with the highest AUPRC among the Wu2026-pretrained host-unseen models at each quantile (Fig. 2A; Fig. 5A-C). For comparison, we also predicted the phages of the held-out hosts using Wu2026-trained models under the both-unseen inference mode to mimic a clinical scenario of directly applying foreign models on local datasets. Model selection at each quantile was likewise based on the AUPRC scores of the Wu2026-trained models in the both-unseen mode (Fig. 2E; Fig. 5A-C).

**Fig. 5:**
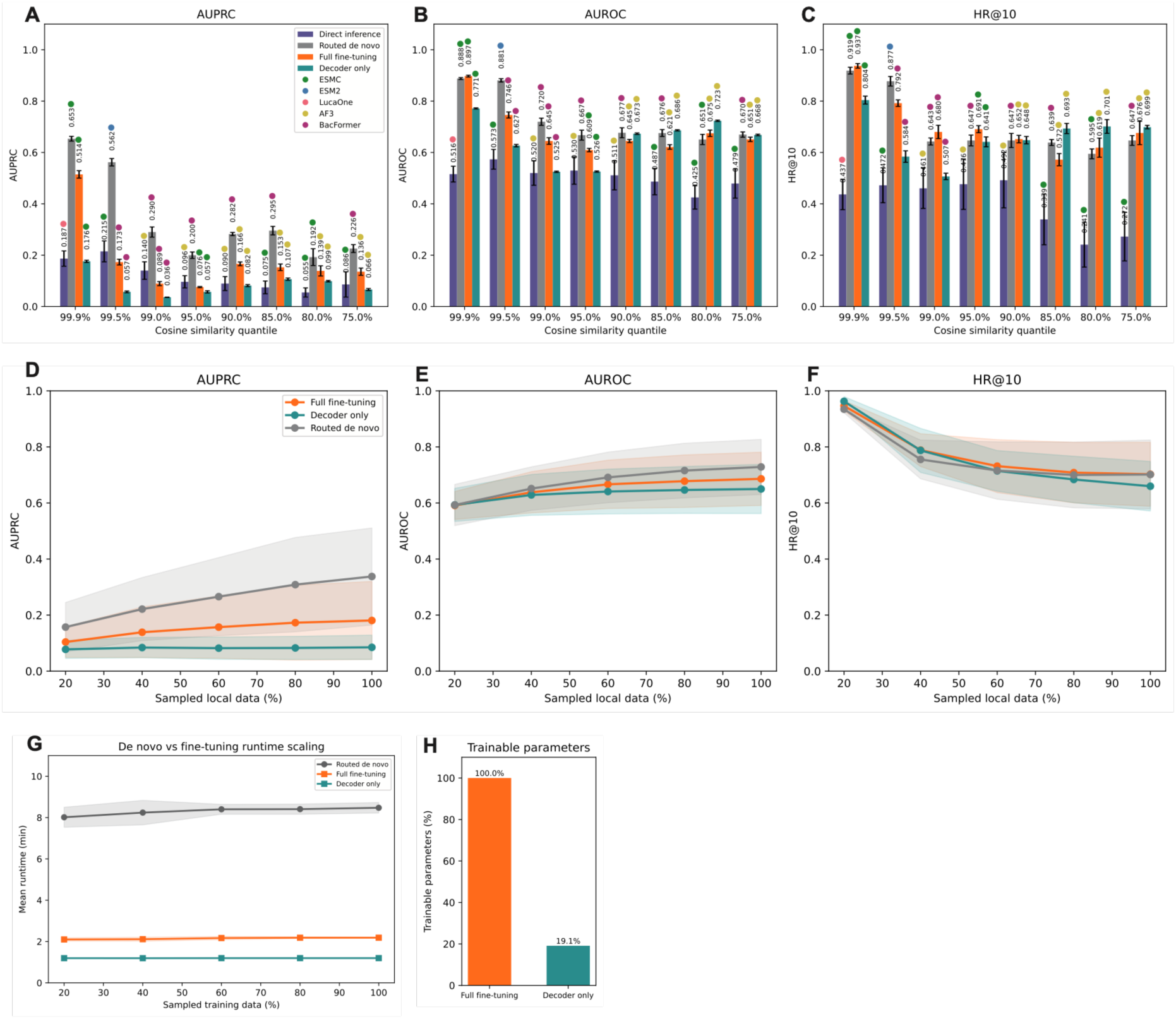
Evaluation of strategies for using the Beamud2023 dataset in phage inference for new hosts. The four strategies were direct inference using the Wu2026-trained models under both-unseen settings, models trained *de novo* on the Beamud2023 dataset under host-unseen settings, and the Wu2026-pretrained models fine-tuned (full and decoder fine-tuning methods) on the Beamud2023 dataset under host-unseen settings. **(A-C)** Bar charts comparing AUPRC, AUROC, and HR@10 across the four strategies at different cosine similarity levels on the entire Beamud2023 dataset. **(D-F)** Performance at the AUPRC-selected best quantile across subsampled interaction fractions, assessed by AUPRC, AUROC, and HR@10 in the full and subsampled Beamud2023 datasets. **(G)** Fine-tuning and *de novo* training runtime across subsampled interaction fractions. **(H)** Numbers of trainable parameters in the full and decoder-only fine-tuning modes. Shaded regions in panels D-G indicate mean ± SD across 10 independent runs per fraction.

Across decreasing similarity quantiles, models trained *de novo* on the local dataset generally performed best, followed by full and decoder-only fine-tuning, whereas direct inference with the Wu2026-trained model in the both-unseen mode generally performed worst (Fig. 5A-C). Specifically, on the whole Beamud2023 dataset, the leading *de novo* models outperformed the fine-tuned models in AUPRC scores at all similarity quantiles (Fig. 5A). In AUROC and HR@10 scores, the leading *de novo* models showed clear advantages at the 99.5%-95% and 99.5% quantiles, respectively; elsewhere performance was comparable to or worse than fine-tuning models (Fig. 5B, C). Full fine-tuning outperformed decoder-only fine-tuning at all host-similarity quantiles in AUPRC, at 99.9%-95.0% in AUROC, and at 99.9%-90% in HR@10; decoder-only fine-tuning performed better at the remaining quantiles. In comparison, direct inference using the Wu2026-trained models on the Beamud2023 dataset showed generally lower performance in AUROC and HR@10 scores than *de novo* and fine-tuning strategies (Fig. 5B, C). In AUPRC scores, direct inference always performed worse than *de novo* training but slightly outperformed both fine-tuning strategies at 99.5% and 95% similarity quantiles, indicating that fine-tuning had limited advantage over direct inference at these two similarity quantiles based on the evaluated settings (Supplementary Dataset 3).

Across all similarity quantiles, the overall performance of the *de novo*-trained models was better (0.338) than the full (0.181) and decoder-only (0.085) fine-tuned models in AUPRC scores (Fig. 5D). The three strategies performed comparably in AUROC and HR@10 scores (Fig. 5E, F). Model evaluation by subsampling the Beamud2023 interaction dataset (from 100% to 20%) showed progressively lower AUPRC for both *de novo*-trained and fully fine-tuned models, suggesting that the performance of *de novo* training and full fine-tuning benefited more from greater interaction coverage than decoder-only fine-tuning, which changed little as interaction-data availability increased (Fig. 5D). In the other two metrics, differences among the three strategies were small regardless of the changes in training data (Fig. 5E, F). Notably, fine-tuned models achieved slightly higher positive retrieval in the top-10 predictions than *de novo* training when the dataset size decreased (Fig. 5F).

Under standardized computational settings on an NVIDIA H100-80 GB GPU, the overall runtime of model training or fine-tuning varied little across dataset sizes (Fig. 5G). Training the different embedding models *de novo* required consistently longer end-to-end model-training wall-clock time (slightly over 8 minutes) than full fine-tuning (slightly over 2 minutes), while decoder-only fine-tuning required the least time (slightly over 1 minute) across 10 runs. Notably, *de novo* training was based on 200 epochs for LOGOCV and 500 epochs for final training, whereas fine-tuning used 50 epochs. Under these settings, full and decoder-only fine-tuning reduced model-training time while updating 100% and 19.1% of the pretrained model parameters, respectively, and retained useful predictive performance (Fig. 5H).

## Discussion

This study presents PhageHost-GNN, a GNN framework that leverages adsorption-stage strain-level specificity between phage and *Klebsiella* strains based on the phage RBPs and bacterial K-locus proteins. Based on the performance of five embedding methods across levels of query novelty, we implemented a novelty-aware model-selection mechanism in the inference procedure. Retrospective benchmarking on host-unseen and both-unseen settings showed that the routed strategy was generally competitive with other approaches in the host-unseen benchmark, while its advantages were clearly observed over fixed embeddings under the both-unseen benchmark. To support model generalization and local adaptability, we developed a pipeline through which users can fine-tune pretrained models on their custom datasets using selected settings and embedding models. The evaluation showed hit ratios comparable to those of *de novo* training, especially for novel *Klebsiella* strains, while using approximately one-quarter of the model-training wall-clock time.

The GNN structure is designed to learn strain-level compatibility patterns. GraphSAGE and GATv2 are suited to link-prediction tasks because GNNs can model biological relationships through convolutions over node features. Furthermore, the GraphSAGE inductive design permits prediction on unseen nodes using a fixed-size sampling technique (local neighborhood), while GATv2 introduces an attention mechanism, assigning weights to neighboring nodes to ensure prioritization of informative neighbors [29–31]. Building on previous GNN applications to phage-host prediction [29, 31], this study uses within-type similarity edges for inductive context propagation and novelty-aware routing across refined phage and host representations, including in the stringent both-unseen setting. [30, 32]. The inductive GraphSAGE design and GATv2 attention mechanism provide a basis for learning strain-level compatibility patterns and making predictions for unseen entities across similarity settings.

The routing strategy of model selection based on the novelty of query entities was used in inference and fine-tuning of our framework, allowing use of the best-performing models for strains at different novelty levels. Multi-representation strategies have previously been explored for phage-host prediction [12, 33–37]. Although previous studies have evaluated prediction performance across different degrees of query novelty or sequence similarity [9, 33], to our knowledge, none have independently quantified phage- and host-side coverage relative to the training data and used coverage-stratified validation performance to hard-select a single embedding model for each query pair. Our model-routing approach was motivated by the complementary performance observed by the five embedding models during LOGOCV evaluation, whereby different embedding models generalized well under different similarity levels (Fig. 2 and 3; Fig. S1 and S2). Therefore, the resulting routed predictions combine the strengths of different embeddings under diverse novelty regimes for adaptive predictions.

Furthermore, the performance comparison among strategies based on direct inference of foreign models, the *de novo* training on local datasets, and the fine-tuning of pretrained foreign models on local datasets examines three options available to a clinical user to match phages in a local library for newly discovered bacterial strains. Direct prediction using a foreign model, trained on datasets geographically distinct from the local library, may result in poor performance depending on the generalization of the model. Our results show that *de novo* training achieved higher AUPRC on custom datasets, while full fine-tuning offered a balance between performance and computational efficiency, reducing model-training time while achieving comparable HR@10 despite updating all model parameters. Furthermore, de novo training achieved stronger classification performance as local dataset size increased, whereas fine-tuning showed advantages in hit ratio. Therefore, we suggest that the three options may be useful under different scenarios, i.e., the availability of local interaction datasets, the degree of phage or host novelty relative to the training entities, and the availability of computational resources (Fig. 5). For instance, direct inference can be applied when users lack or have limited interaction datasets, especially when new phages or host strains are similar to the training datasets. Fine-tuning may be a suitable option when local interaction datasets and computational resources are limited, while *de novo* training can be leveraged when sufficient representative target-domain data become available for comprehensive learning under the new biological context. However, these conclusions are drawn from evaluations on the Beamud2023 dataset, and additional prospective validations on independent datasets are required before broader application.

In conclusion, PhageHost-GNN integrates RBP and K-locus representations with novelty-aware graph learning to rank candidate *Klebsiella* phage–host pairs across host-unseen, phage-unseen, and both-unseen settings. Retrospective benchmarks and one cross-regional adaptation experiment show that the framework can enrich experimentally verified interactions among top-ranked candidates. These findings support its use as a prescreening tool to prioritize wet-lab testing. Prospective multisite evaluation is required to determine whether it reduces screening time or cost in phage-therapy practice. Because RBPs and K-locus proteins were prespecified model inputs, the results are consistent with—but do not independently establish—their role in host-range specificity.

### Limitations of the study

Despite the promising predictive performance, we acknowledge several limitations. For instance, interactions that do not use the capsule or K-antigen as the primary receptor may be missed because the model is restricted to RBP and K-locus inputs. Additionally, receptor-mediated adsorption is only the first step of productive infection. Post-adsorption host defenses and intracellular compatibility can block genome entry, replication, assembly, or lysis even after binding, and these determinants are not represented by the current interaction mechanism [38]. In addition, kNN auxiliary edges may introduce similarity bias for divergent entities; therefore, future evaluation should quantify these effects with denser interaction matrices and independent controls. Although the model was evaluated retrospectively on experimentally validated datasets, prospective evaluation is required to assess its generalizability in practice.

## Materials and Methods

### Dataset preparation

This study used multiple datasets for training and evaluation, as shown in Table 1. The experimentally verified interaction networks in these datasets were obtained from public repositories from previous studies [1, 8, 11, 28]. The genome sequences of the phage and host strains were either obtained from the data repositories cited in the original studies or the NCBI database using the reported accession numbers when sequences were not available in the original repositories.

### Sequence data collection and processing

For the Boeckaerts2024 dataset, phage RBPs and capsule synthesis locus (K-locus) proteins of the *Klebsiella* hosts were obtained from the online deposit of the original study [11]. For the other datasets, genes were identified from phage genome sequences using PHANOTATE [39] and translated in silico into proteins; phage RBPs were then detected using PhageRBPdetection v4 [5], with RBPs shorter than 200 or longer than 1,500 amino acids excluded to match the expected lengths defined by Latka et al. [40]. *Klebsiella* genome sequences were processed using Kaptive v3.0 [41–43] to identify the best-matching K-locus type from which K-locus protein sequences were extracted for subsequent processing.

### Protein structure representations

Phage RBPs and bacterial K-locus proteins were submitted to AF3 [23, 24] for protein-structure prediction using the AF3 database (custom 2024-11-16, PDB:2022-09-28) and default parameters on multiple NVIDIA H100 80 GB GPUs (Fig. 1A). Structural representations were extracted from AF3 CIF structures using the ESM-IF1 encoder [26], which processes protein backbone atoms (N, Cα, C) from CIF structures based on the geometric vector perceptrons (GVP) and converts them into structural representations.

### Protein language model representations

Numerical vector representations of all phage RBPs and host K-locus protein sequences were generated via pretrained biological foundation models, namely LucaOne (GPLM v2.0, step 5.6M) [22], ESM2 (t33_650M_UR50D) [21], ESMC (600M, 2024_12_v0 checkpoint) [20], and Bacformer (bacformer-masked-complete-genomes checkpoint) [25], which were implemented on their respective standard architectures on a single NVIDIA H100 80 GB GPU in dedicated Conda environments. These models take single protein sequences as input and produce unique, fixed-length numerical vector representations based on the specific model configurations, including 2560-, 1280-, 1152-, and 480-dimensional representations for the LucaOne, ESM2, ESMC, and BacFormer, respectively, while AF3 produced 512-dimensional embeddings extracted via the ESM-IF1 encoder (Fig. 1A).

### Multi-instance feature aggregation

Multi-instance structural and LLM protein vectors from phages and bacterial hosts were aggregated using a column-wise mean, i.e., 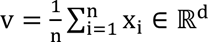, where *n* is the total number of embeddings per organism, and the embedding of protein i is represented by *x_i_* ∈ ℝ*^d^*. This approach summarizes multiple protein instances while preserving the embedding dimension; the resulting vector represents each phage or bacterium and serves as the input node feature for GNN representation learning.

### Classification model training

Column-wise aggregated representations served as node features for model training over a fixed number of epochs: 200 epochs for LOGOCV and 500 epochs for final training from random initialization on the full training dataset, using a learning rate of 0.001 and a weight decay of 0.005. The model was implemented in a heterogeneous GNN pipeline using Python v3.9.23 and PyTorch v2.2.2 [44] and trained on a single NVIDIA H100 80 GB GPU using modality-specific parameter configurations. Phage and host node features were separately projected into 256-dimensional hidden representations, and stacked SAGEConv and GATv2Conv layers were employed for local feature aggregation and multi-head attention (two heads), respectively, with residual updates that preserve the initial features to facilitate type-specific message passing and obtain enriched representations. An MLP edge decoder scores and ranks candidate pairs using adjacent phage-host embeddings. During training, the overall GNN architecture remained consistent across modalities, with fixed primary training hyperparameters to support fair performance comparisons; input dimensions varied across model configurations (Supplementary Tables S7–S8).

### Graph convolution and interaction prediction

The proposed multimodal GNN framework for predicting *Klebsiella* phage-host interactions was implemented using the GraphSAGE (SAmple and aggreGaTE) and Graph Attention Network (GATv2) algorithms (Fig. 1B), leveraging LLM and structural embeddings as node features for feature aggregation and dynamic multi-head attention, respectively. We define a heterogeneous network *G = (V, E)*, where *V* comprises phage and host nodes, while *E* denotes the set of experimentally validated interactions in the matrix M ∈ {0,1}^m×n^, where *m* and *n* are the number of phages and host strains, respectively. M_ph_ = 1 for experimentally validated positive interactions and 0 for experimentally validated negative interactions, whereby phage p ∈ {1, …, m} and host h ∈ {1, …, n}. Unobserved pairs were excluded from the training and evaluation datasets. Each node *i* contains a structural or LLM embedding 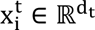, with adjacent node embeddings facilitating the computation of its interaction scores via an MLP decoder.

Message passing was conducted through directed phage-phage and host-host cosine-similarity edges, connecting each node to its top-10 k-nearest neighbours (kNN), while experimentally verified interactions were employed for supervision by the decoder. kNN graph reconstruction was independently conducted at every LOGOCV fold after node representations were standardized by mean and standard deviation based on the corresponding training entities. The training graph only contained training entities, which permitted two-way message passing, whereas the held-out test nodes were limited to receiving messages from nodes in the training graph to minimize the risk of data leakage for the test nodes while allowing propagation from the training graph. Experimentally validated phage-host interactions were used for supervision by the model decoder.

Inductive node representations were initially obtained using GraphSAGE, a GNN algorithm based on neighbour sampling and feature aggregation [29], as shown in Equation 1.

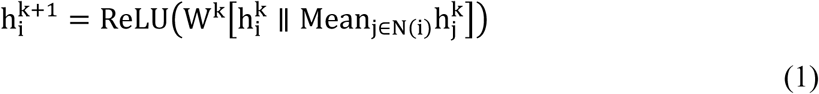

Here, *W^k^*denotes the learned weights in layer 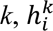 denotes the initial representation of node *i, N(i)* is the set of sampled neighboring nodes *j* used to compute the enriched representation 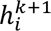 of the subject node *i*, and ∥ denotes concatenation.

Multi-head attention is employed via GATv2, with two attention heads (H) after each SAGEConv layer to weight the contributions of neighboring nodes [45, 46]. As a result, the attention coefficients for a given edge *e_i_*_j_and attention head h ∈ H are computed from learned compatibility scores and normalized over the neighborhood *j* ∈ *N*(*i*), before the weighted neighborhood features are averaged across the multiple attention heads for the final node embedding, as shown in Equation 2.

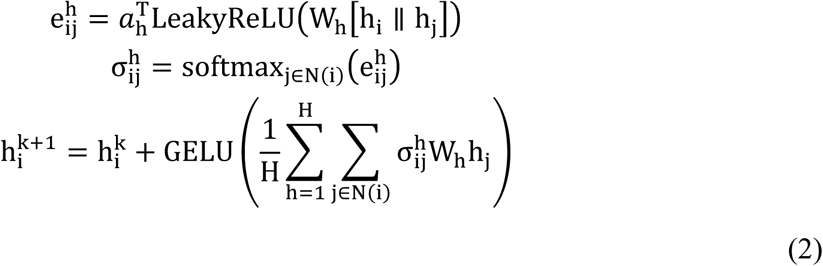

Here, 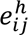 denotes the dynamic and unnormalized attention coefficients of the subject node *i* and its neighbouring node *j* in attention head *h*, *a*_ℎ_ are learnable attention vectors that convert projected node features into compatibility scores to determine the relevance of a given neighbourhood, *W*_ℎ_ is the learnable feature-transformation weight matrix, while 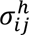 are normalized attention coefficients obtained via the softmax function. 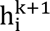 represents the final updated node representation via multi-head averaging, with a residual update and GELU activation to stabilize feature propagation and minimize the risk of over-smoothing, while preserving functional and evolutionary signals from the respective modalities. Ultimately, an MLP edge decoder operates on the concatenated 256-dimensional adjacent GNN-enriched embeddings, their elementwise products, and absolute differences, allowing the model to capture compatibility and embedding differences and to produce projected prediction scores from enhanced GNN representations for ranking and experimental validation.

### Inference

During inference, new hosts were connected to the 10 most similar counterparts in the training graph, which comprised all training phages and hosts. Consistent with a previous study [11], each bacterial isolate was scored against a subset of the training phages by the respective quantile-assigned models, generating ranked predictions. In the both-unseen inference mode, each novel phage-host pair is scored by the respective quantile-assigned models, after which the model predictions can be evaluated as a phage or host ranking system. For binary classification on independent inference datasets, decision thresholds were generated for the respective model quantiles using the training cross-validation precision-recall curve. Precision (P) and recall (R) were computed from the pooled predictions for all candidate thresholds, and the F1-score was calculated for each threshold as the harmonic mean of precision and recall, i.e., F1 = 2PR/(P + R) and the threshold that maximized the F1-score was selected as the optimal threshold (Supplementary Table S11).

### *In silico* model performance evaluation

*In silico* performance was evaluated across three LOGOCV training configurations: phage-unseen, host-unseen, and both-unseen, simulated by iteratively holding out a single phage or host group as the test set with its corresponding training-set interactions, and predicting the most active candidates. For both-unseen configurations, one phage cluster and one host cluster were simultaneously held out and paired exhaustively by Cartesian combinations as a test set, while the remaining clusters were used for training. Held-out pairs with at least one observed interaction were retained, and the model performance was evaluated on labeled pairs within held-out clusters. Furthermore, all supervision edges associated with the held-out entities were excluded from training to minimize data leakage. Groups were generated via greedy clustering, where unassigned entities from the similarity matrix were clustered with remaining entities above the cutoff using empirically defined cutoffs, with identical entities clustered at the highest cutoff. Embedding cosine-similarity clustering using defined quantiles (99.9% to 75%) was employed on the three training configurations, with a single threshold computed per quantile using off-diagonal similarities, and every entity pair was compared against the threshold to generate quantile-based clusters due to the existing differences in modality-specific feature distributions (Supplementary Table S10).

The model was evaluated using the AUROC, AUPRC, and HR@K metrics calculated from all held-out predictions pooled across all LOGOCV folds (micro-pooling). Model predictions were ranked for every eligible query, and a hit was recorded when at least one positive interaction existed within the top-K predictions. The same process was applied across different similarity groups, with the query HR@K macro-averaged on all eligible entities, after excluding entities without observed positive interactions. The final HR@K therefore represents the proportion of eligible query entities for which at least one positive interaction was retrieved among the top-K predictions.

### Model selection and retrospective evaluation

In the host-unseen evaluation, host strains were assigned to the respective quantiles based on their similarity to the hosts in the training dataset, while in the both-unseen evaluation, each phage and host were assigned to their respective quantiles, and the lower of the two was selected as the supporting quantile (Algorithm 1). Highly novel query entities that were not supported by any similarity quantile across all embedding models were regarded as out-of-distribution entities and evaluated using the lowest similarity quantile models (75%) as an extrapolation fallback. However, none of the queries in the current evaluation required this fallback because all entities were supported by at least one embedding model. For inference of each new host, a model was selected based on the highest training AUPRC at the assigned quantile, with a similar approach employed in the both-unseen evaluation, in which the highest AUPRC model was selected for the lowest phage-host pair quantile. Retrospective evaluation was then conducted using the three performance evaluation metrics against the ground truth of the *in vitro* datasets.

#### Algorithm 1

Both-unseen similarity-guided model routing

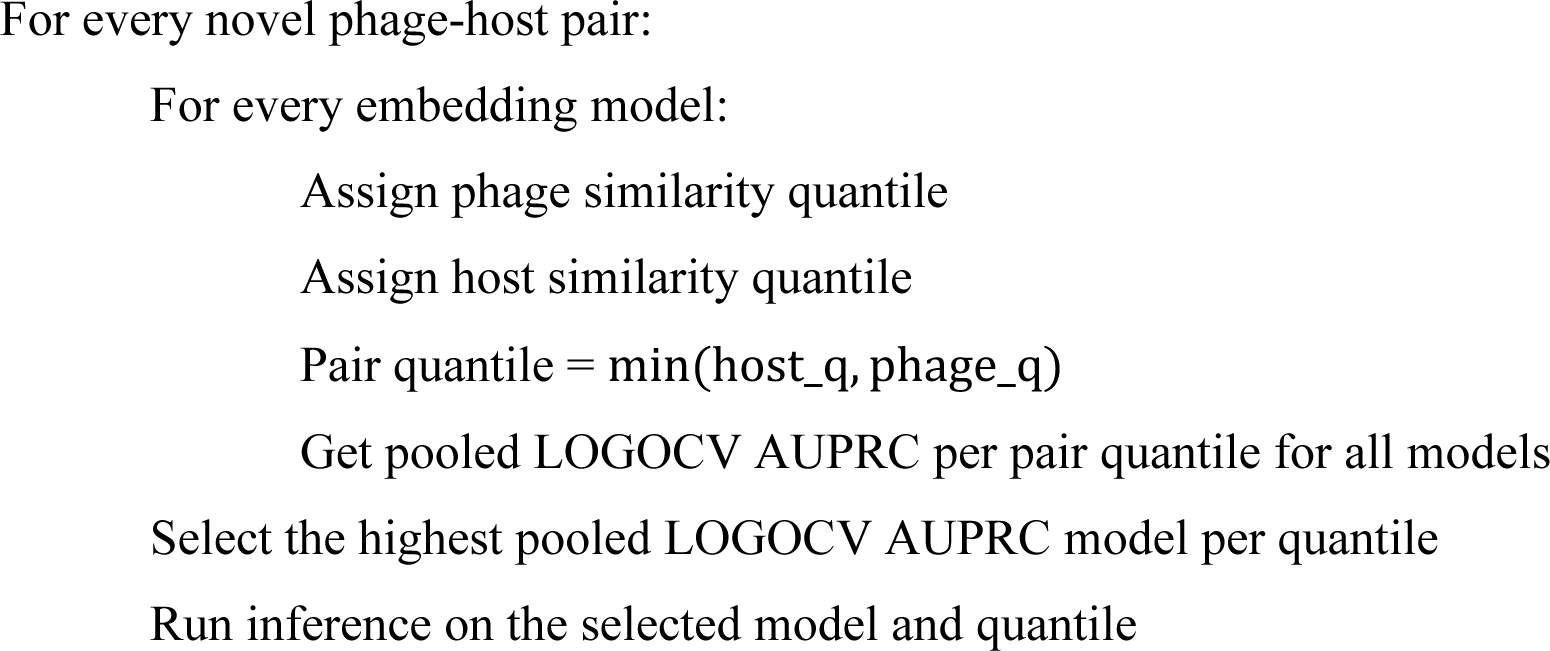

### Local adaptation

A local adaptation pipeline was evaluated using full fine-tuning and decoder-only PEFT, adapting Wu2026-pretrained models to the Beamud2023 dataset in the host-unseen mode (Fig. 1E). Pretrained models were fine-tuned for fewer epochs using their original training configurations. Bacterial strains were clustered by embedding cosine similarity across quantiles from 75% to 99.9% for LOGOCV training, with one group held out for testing and the remaining groups used for adaptation based on the pretrained model with the highest AUPRC per quantile (Fig. 2A). For comparison, *de novo* training was conducted on the remaining groups and then tested on the held-out group under the host-unseen mode using the five embedding methods. In addition, direct inference based on the Wu2026-pretrained models on the held-out group was conducted at each quantile in the both-unseen mode using the best-performing embedding method shown in Fig. 2E. Fine-tuning and de novo training were conducted on an NVIDIA H100 80 GB GPU. Adaptation was evaluated on 20–100% stratified samples of the interaction dataset to examine performance across different levels of interaction data coverage and compare full fine-tuning, decoder-only fine-tuning, and de novo training.

Each setting was repeated with 10 random seeds, and the performance summary was reported as mean ± SD (Supplementary Table S9).

## Grants

This study was supported by the National Key R&D Program of China (2024YFE0198900 to K.Z.), the National Natural Science Foundation of China (No. 42376113 to L.F.), and the Shenzhen Key Laboratory of Marine Archaea Geo-Omics, Southern University of Science and Technology (SYSPG20241211173725010 to L.F.).

## Ethics declaration

Not applicable.

## Data availability

Datasets are publicly available via GitHub: https://github.com/LoMI-SUSTech/PhageHost-GNN

## Code availability

Model implementation code is publicly available via GitHub: https://github.com/LoMI-SUSTech/PhageHost-GNN

Protein sequences were processed using:

PHANOTATE (https://github.com/deprekate/PHANOTATE),

PhageRBPdetection v4.0 (https://github.com/dimiboeckaerts/PhageRBPdetection),

Kaptive v3.0 (https://github.com/klebgenomics/Kaptive)

Protein embeddings were computed using:

ESMC (https://github.com/Biohub/esm),

ESM2 (https://github.com/facebookresearch/ESM),

LucaOne (https://github.com/LucaOne/LucaOne),

BacFormer (https://github.com/macwiatrak/Bacformer),

AlphaFold 3 (https://github.com/google-deepmind/alphafold3).

## Author contributions

Conceptualization and methodology: B.M.M., L.F.; Writing - original draft: B.M.M.; Writing - review and editing: B.M.M., K.Z., L.F.; Software: B.M.M.; Data curation: B.M.M., K.Q.; Formal analysis: B.M.M.; Supervision: K.Z., L.F.; Project administration: L.F.; Resources: K.Z., L.F.

## Acknowledgements

We thank Jie Feng from Institute of Microbiology, Chinese Academy of Sciences, Rui Zhang from Shenzhen University, and Yanqing Hu from Southern University of Science and Technology for their advice in data interpretation and model selection. Computation in this study was supported by the Center for Computational Science and Engineering at the Southern University of Science and Technology.

