## Supplementary Information for "PhageHost-GNN: novelty-aware and locally adaptable strain-level *Klebsiella*-phage prediction"

### Supplementary Figures

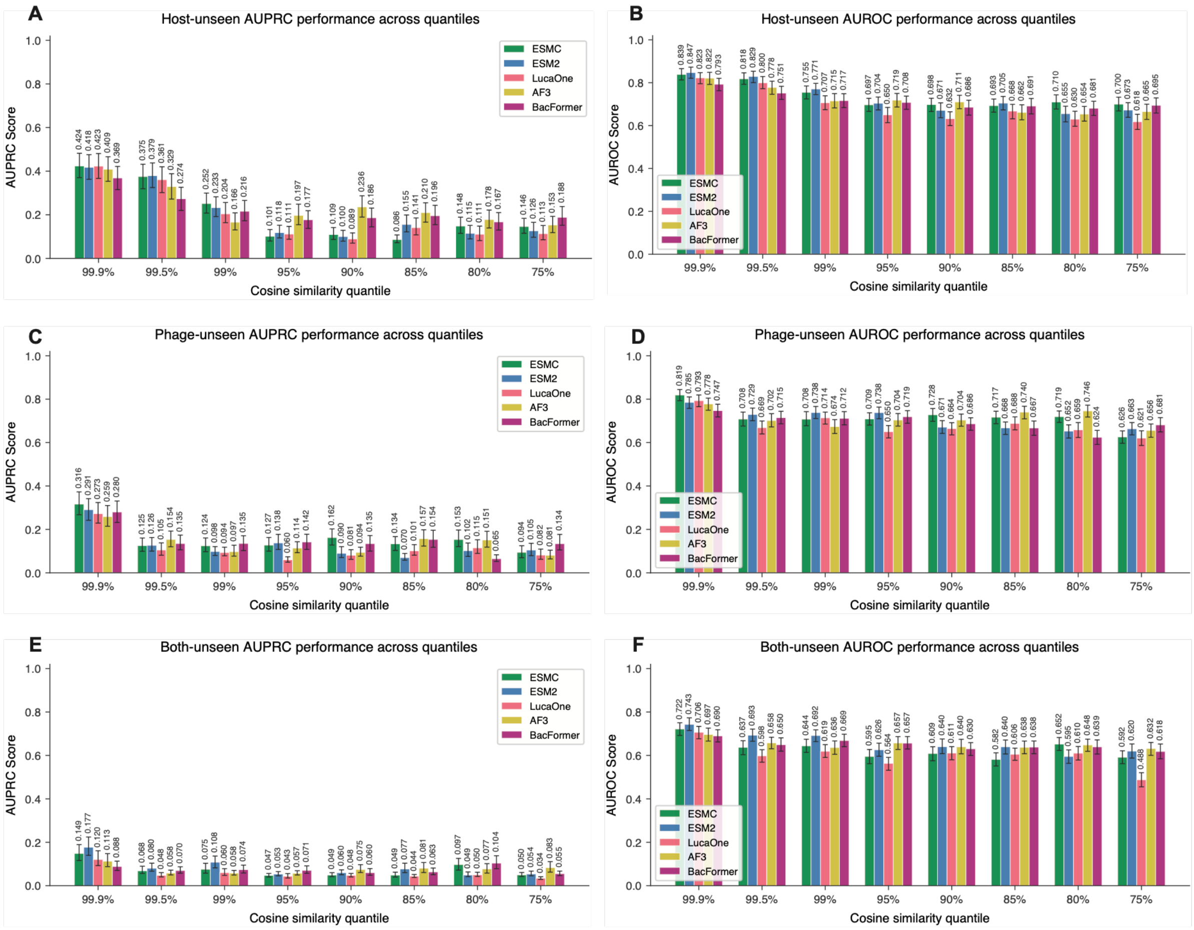


**Fig. S1: AUPRC and AUROC performance of the PhageHost-GNN models on different embeddings trained on the Boeckaerts2024 dataset on hosts/phages of different novelties.** LOGOCV was conducted on the Boeckaerts2024 dataset at each cosine similarity clustering at each cosine-similarity threshold. The held-out cluster was used for inference, while the remaining clusters were used for model training. (A, B), (C, D), (E, F) show the AUPRC and AUROC scores obtained under the host-unseen, phage-unseen, and both-unseen inference modes, respectively.

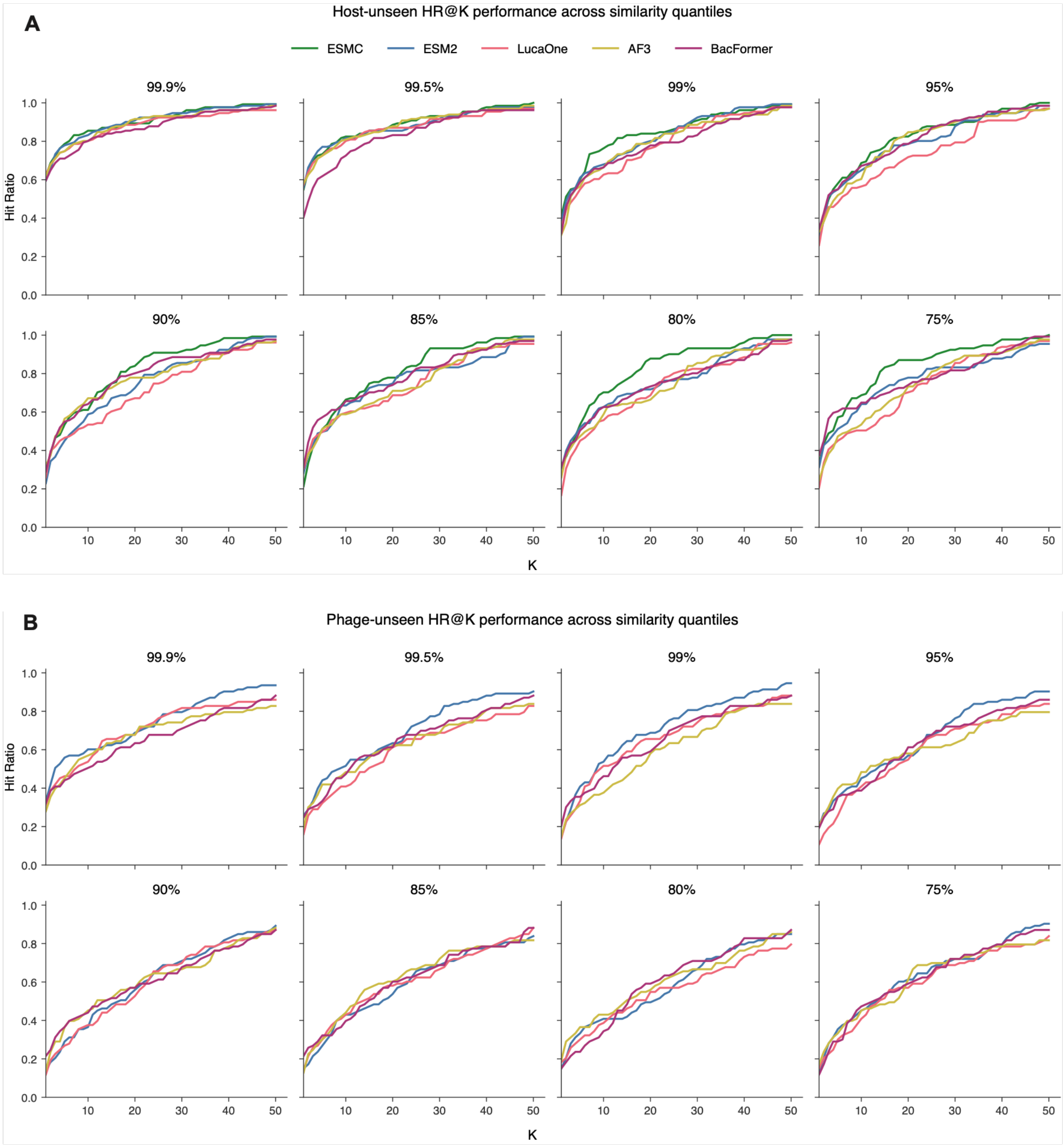


**Fig. S2: HR@1-50 curves for the PhageHost-GNN models of different embeddings trained on the Boeckaerts2024 dataset on hosts/phages of different novelties under host-unseen (A) and phage-unseen (B) inference modes.** LOGOCV was conducted on the Boeckaerts2024 dataset at each cosine-similarity threshold. The held-out cluster was used for inference, while the remaining clusters were used for model training.

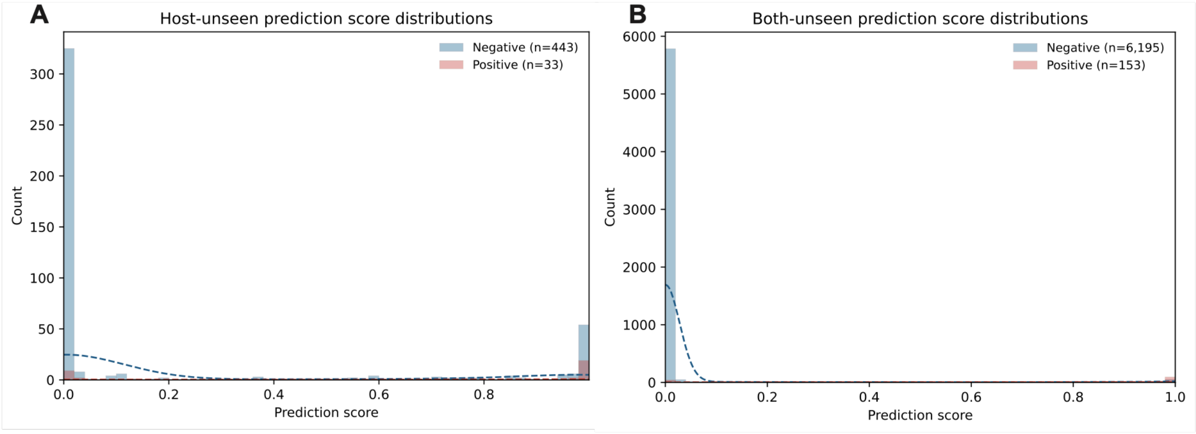


**Fig. S3: Prediction-score distributions in the host-unseen setting for the Boeckaerts2024 (A) in vitro datasets and in the both-unseen setting for the Beamud2023 dataset (B).**

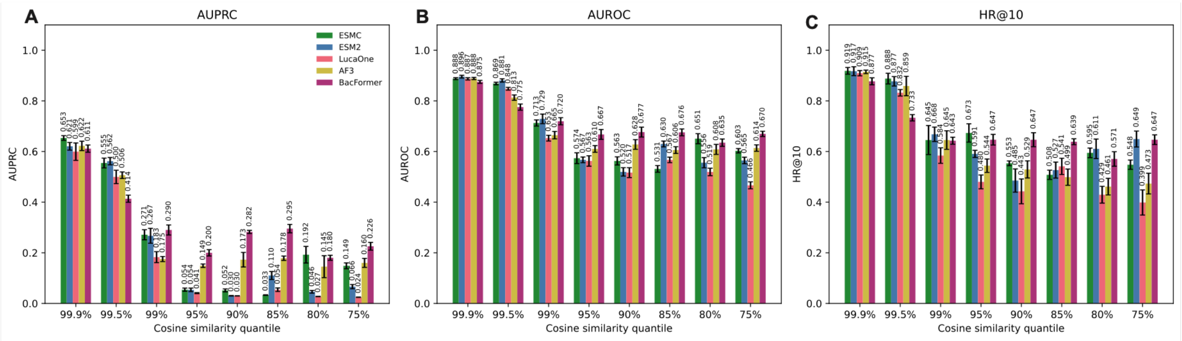


**Figure S4:** De novo host-unseen model performance on the Beamud2023 dataset showing AUPRC, AUROC, and HR@10 for the respective models across decreasing host similarities.
